# Molecular-scale 3D visualisation of the cardiac ryanodine receptor clusters and the molecular-scale fraying of dyads

**DOI:** 10.1101/2021.10.21.462365

**Authors:** Thomas M. D. Sheard, Miriam E. Hurley, Andrew J. Smith, John Colyer, Ed White, Izzy Jayasinghe

## Abstract

Clusters of ryanodine receptor calcium channels (RyRs) form the primary molecular machinery in cardiomyocytes. Various adaptations of super-resolution microscopy have revealed intricate details of the structure, molecular composition and locations of these couplons. However, most optical super-resolution techniques lack the capacity for three-dimensional (3D) visualisation. Enhanced Expansion Microscopy (EExM) offers resolution (in-plane and axially) sufficient to spatially resolve individual proteins within peripheral couplons and within dyads located in the interior. We have combined immunocytochemistry and immunohistochemistry variations of EExM with 3D visualisation to examine the complex topologies, geometries and molecular sub-domains within RyR clusters. We observed that peripheral couplons exhibit variable co-clustering ratios and patterns between RyR and the structural protein, junctophilin-2 (JPH2). Dyads possessed sub-domains of JPH2 which occupied the central regions of the RyR cluster, whilst the poles were typically devoid of JPH2 and broader, and likely specialise in turnover and remodelling of the cluster. In right ventricular myocytes from rats with monocrotaline-induced right ventricular failure, we observed hallmarks of RyR cluster fragmentation accompanied by similar fragmentations of the JPH2 sub-domains. We hypothesise that the frayed morphology of RyRs in close proximity to fragmented JPH2 structural sub-domains may form the primordial foci of RyR mobilisation and dyad remodelling.

## Introduction

Myocardial contraction is enabled through synchronous calcium (Ca^2+^) release within each cardiomyocyte. Ryanodine receptor (RyR) Ca^2+^ channels, clustered on the termini of the sarcoplasmic reticulum (SR) and activated by Ca^2+^-induced Ca^2+^ release (CICR), form the primary route of this release of Ca^2+^ into the myoplasm. Spatiotemporal synchrony of the elementary events of Ca^2+^ release (Ca^2+^ sparks (1)) is paramount to achieving the cytoplasmic concentrations required for activating a fast and forceful muscle contraction. Vital to ensuring the regenerative nature and synchrony of Ca^2+^ release are a number of key determinants of the local control of RyRs. These include sub-micron-scale co-localisation of the primary trigger, the L-type Ca^2+^ channel (LTCC), and other components of the excitation-contraction (EC) coupling machinery (2), Ca^2+^-dependent inactivation of RyR, the [Ca^2+^] in the SR lumen, post-translational modifications of RyR, the availability and binding of modulators such as FKBP variants and junctophilin-2 (JPH2), and the geometry and distribution of the RyR clusters (see review (3)).

The clustering properties of the cardiac RyRs, in terms of cluster size and the intra- and inter-cluster spacing, are key determinants of cluster excitability (4,5) and the variability of the Ca^2+^ spark properties (6). Observations of closely-arranged groups of clusters have further led to the hypothesis of functionally-coupled ‘super clusters’, in which neighbouring clusters may be co-activated by diffusing Ca^2+^ (7). Additionally, the non-uniform self-assembly of RyR arrays within the dyads (8,9), the size of the dyad (2,10) and the geometry of the dyad linked to the shapes and sizes of the t-tubules (4), are key determinants of the local control of the RyRs within. Unsurprisingly, many of these structural proteins also form a critical part of the primordial dyads which establish the contractility of developing cardiomyocytes (11,12).

Regulating the organisation of RyRs and the EC coupling machinery are several key structural molecules such as JPH2 and amphiphysin-2 (BIN1). JPH2, which also works as a direct modulator of RyR, physically tethers the SR and the sarcolemma (13) and maintains the shape of the t-tubule and the size of the RyR cluster (14,15). Loss or downregulation of dyad-related structural proteins has been linked to the aetiology of the maladaptive remodelling of the t-tubules and dyad structures, and concurrently, the dysregulation of the intrinsic Ca^2+^-handling in a range of cardiac pathologies (16). We (5) and others (6,17) have shown previously that this remodelling extends to the RyR organisation at both dyads and sub-sarcolemmal (peripheral) couplons.

Many of the newer insights into sub-cellular remodelling and molecular-scale organisation of RyR have come from advanced optical and electron microscopy techniques. Optical super-resolution microscopies (known best by acronyms such as STED, dSTORM, DNA-PAINT) have led the way in visualising both cellular compartments and precise molecular targets such as RyRs with nanoscale resolution (< 250 nm; see review: (18)). Tomographic electron microscopies (EM) have advanced our view of the three-dimensional (3D) complexities of couplon and dyad structures (8,9,19,20), whilst advanced EM-based staining techniques have also enabled *in situ* mapping of macromolecules such as RyR (8). *In situ* molecular counting approaches have been enabled by the newer optical super-resolution techniques, particularly as the strategies to standardise single-molecule photochemistry, the sustainability of the probes and resolution have continued to improve. For example, DNA-PAINT, offering ≤ 10 nm resolution and a target counting algorithm (qPAINT), has become instrumental in quantifying both RyR cluster sizes and the natural heterogeneity in the co-clustering ratio between RyR and JPH2 (21). The recent application of enhanced expansion microscopy (EExM), a swellable hydrogel approach to obtain resolution of ∼ 15 nm, has allowed individual channels to be visualised and counted within clusters located deeper in the myocytes, both in healthy and failing hearts (5). However, a major advantage of EExM has been the far superior axial resolution (≤ 35 nm) and imaging depth compared to the molecular-scale imaging protocols such as DNA-PAINT, which currently set the benchmark in optically resolving single dyad targets.

In this article, we demonstrate the utility of EExM variants (both 10x and 4x; (5)) in visualising the topologies and geometries of RyR arrays *in situ*. We explore the use of both immunocytochemistry and immunohistochemistry to gain different perspectives of the RyR cluster and t-tubule geometries. By combining 3D visualisation with molecular counting of RyR and JPH2, we report the nanoscale dyad remodelling which accompanies the dissipated or fragmented morphology of RyR clusters observed to-date in cardiac pathology in monocrotaline (MCT)-induced right ventricular (RV) heart failure.

## Results

### Characterization of 3D resolution attainable with EExM

10x EExM is an imaging protocol which combines the ∼1000-fold inflation of a hydrogel-based fluorescent imprint of the sample, based on a protocol called “X10 ExM” (24), with a further 2-fold resolution improvement afforded by Airyscan over regular confocal microscopy (5). Figure 1A illustrates an exemplary confocal micrograph of immunofluorescence labelling of RyR2 in a rat ventricular myocyte. With 10x EExM, not only can the banded RyR staining morphology be resolved to be domains with intricate and varied shapes (Figure 1B), but the nanoscale punctate labelling densities, which represent individual RyRs (21), are also clearly observable. Maximum-intensity z-projection of one of these clusters, colour-coded for depth (Figure 1C), reveals the intrinsic ability of 10x EExM to not only resolve individual RyRs, but also detect them at a 3D scale well below the ‘diffraction limit’.

**Figure 1.**
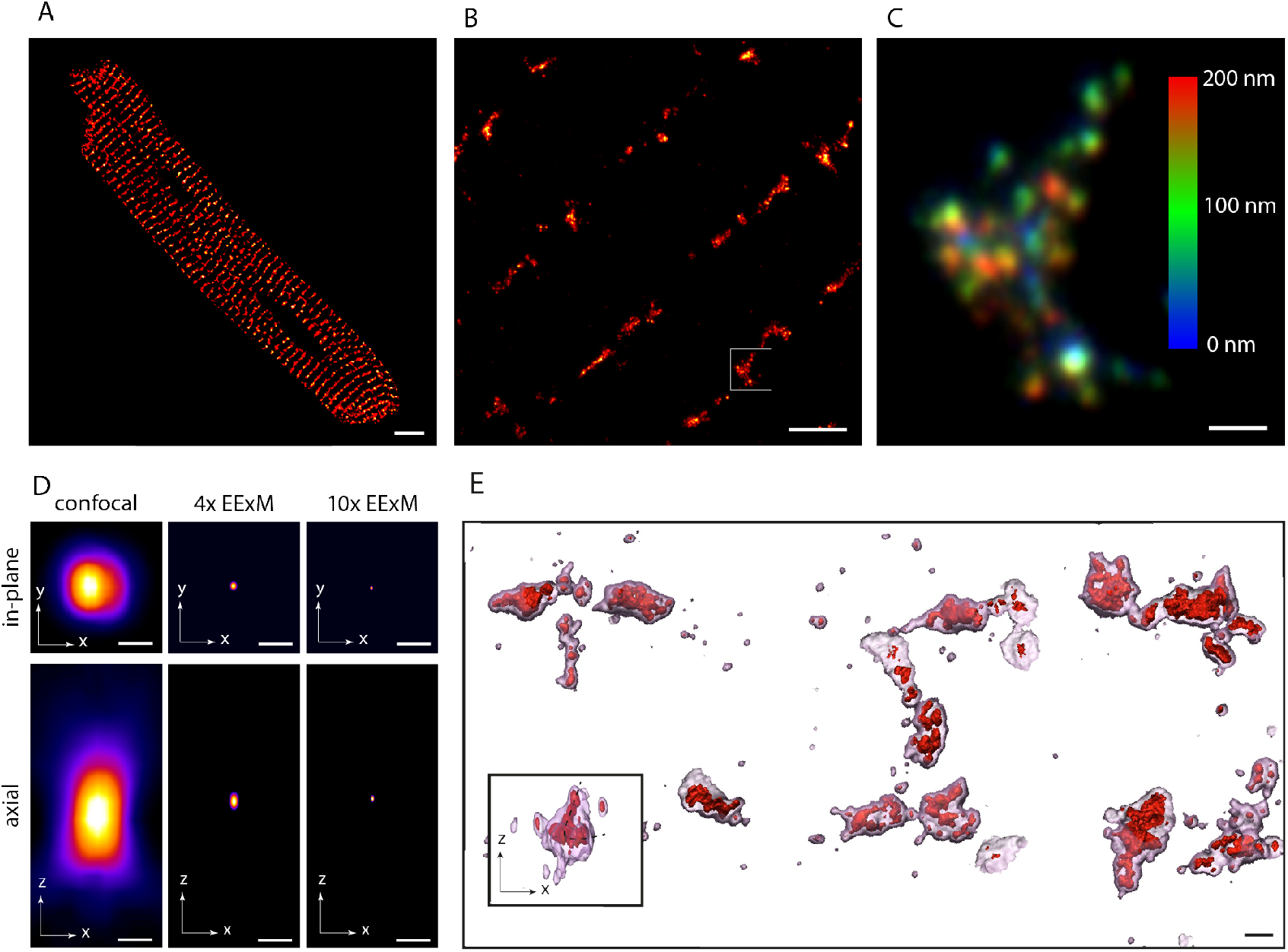
Three-dimensional complexity of ryanodine receptor clusters visualised with EExM. **(A)** A typical 2D Airyscan image of RyR2 immunofluorescence labelling in a rat ventricular myocyte. **(B)** A 2D Airyscan image of a region in a similar cell, acquired with the 10x EExM protocol. **(C)** Magnified and projected view of the region of interest in panel B, colour-coded for axial depth. Colour bar indicates depth in nanometres. **(D)** To-scale comparison of the effective point spread functions of confocal, 4x EExM and 10x EExM, shown in the in-plane (upper) and axial dimensions (lower). **(E)** Isosurface rendered 3D reconstruction of two adjacent rows of RyR, visualized with 10x EExM (red) and a resolution equivalent to 4x EExM (transparent pink), illustrating the superior resolution and better-resolved curvatures of the RyR clusters with the former; inset shows the x-z view of an exemplary cluster whose curved topology in the z-dimension (dashed line) was better-resolved with 10x EExM than with 4x EExM. Scale bars: A: 5 μm; B: 1 μm; C, D & E: 100 nm.

This improvement in both in-plane and axial resolution is a direct result of the effective shrinking of the confocal point-spread function by a factor of ∼ 20 in each of the three dimensions (Figure 1D). Even 4x EExM (which uses the widely-used pro-ExM protocol; (26)) offers an 8-fold resolution improvement over the confocal PSF (see simulations in Supplementary Figure S5 of (5)). When imaging the 3D topologies of the RyR clusters, 4x EExM therefore produces sharply-defined cluster shapes (transparent pink isosurface in Figure 1E), qualitatively similar to the morphology seen with 3D dSTORM (28). With 10x EExM however, we were able to observe the cluster sub-structures, including curved or folded arrays of RyR (solid red isosurface in Figure 1E), which were not observed in previous STORM studies of RyRs located in the cell interior (6,29,30). With an axial resolution of ∼ 35 nm, 10x EExM was also capable of resolving curvatures of the RyR clusters which extended in the z-dimension (inset of Figure 1E).

### Visualising transverse and longitudinal perspectives of dyads

We and others have previously shown that re-orienting myocytes to scan the RyR clusters in transverse view allows a superior view of the network structure of the dyads and t-tubules compared to the conventional approach (imaging myocytes in longitudinal orientation; (31,32)). By combining 4x EExM with tissue immunohistochemistry, we examined myocytes that were physically sectioned in the transverse plane and visualised these structures at an in-plane resolution < 40 nm and an axial resolution < 90 nm (Figure 2A). Examination of a zoomed-in region of the 4x EExM image of RyR showed clusters similar in outline to previous 2D and 3D dSTORM images (28,29), however the sub-structures of the clusters was heterogeneous and punctate (Figure 2B). This view was ideal for examining the network of t-tubules which extend across the cell’s z-line and optically resolve the tessellation of individual RyR clusters around the tubules. 3D isosurface rendering of the t-tubules illustrates the intricate nanoscale topologies of the t-tubular network (cyan in Figure 2C) whilst a lightly-smoothed rendering of the RyR labelling (red) illustrates the curvatures of the RyR arrays around the local t-tubule geometry. Depth-encoded coloured isosurface rendering of the smoothed RyR image further illustrates the intricate curvatures of the RyR clusters, which were previously not visualised with optical microscopy methods (Figure 2D).

**Figure 2.**
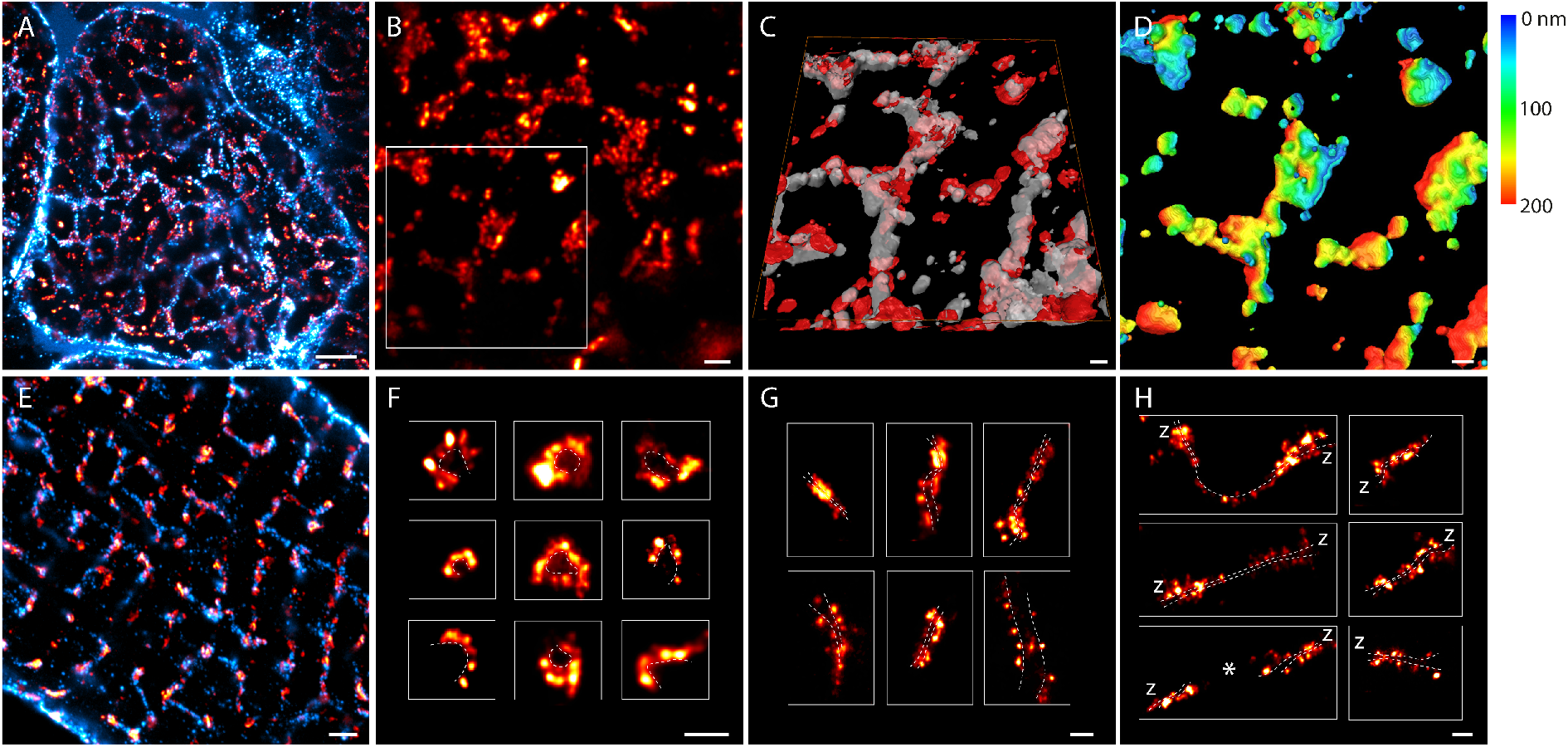
Transverse and longitudinal views of the organisation and geometries of RyR clusters. Transverse view of a ventricular myocyte within a cryosection of right ventricular tissue, labelled for RyR2 (red) and t-tubules (cyan) and imaged with 4x EExM, illustrated by a maximal intensity projection near a single z-line. **(B)** RyR labelling in a 2D, transverse view shows a highly punctate morphology. **(C)** Isosurface rendering of the RyR (red) and t-tubule (cyan) channels illustrate that when imaged in transverse view, even 4x EExM can reveal the topological complexities of the RyR clusters. **(D)** These topologies are shown more directly with an axial depth-encoded (scale shown in nanometres) colouring of a smoothed isosurface rendering of the RyR volume. **(E)** 10x EExM of enzymatically isolated myocytes are more suitable for visualising RyR (red) and t-tubules (cyan) throughout the cell interior in longitudinal view. In this view, we can identify RyR clusters arranged in varying orientations relative to the image plane: **(F)** visualised in “end-on” orientation, **(G)** clusters located at the z-lines, visualised in “side-on” orientation and **(H)** clusters observed to be extending longitudinally or diagonally between z-lines, visualised similarly in “side-on” orientation. Asterisk indicates an example of consecutive longitudinal clusters located on a single longitudinal tubule. As a guide, the approximate location of the respective t-tubule membrane has been indicated with the dashed lines. Scale bars: A & E: 1 μm; B, F, G & H: 250 nm; C & D: 100 nm.

Whilst 10x EExM is currently not compatible with myocardial cryosections (due to the need for more thorough digestion enabling increased expansion factor) it is still highly suited for imaging the RyR clusters located deeper in the interior of enzymatically-isolated myocytes. Typically, imaging of these samples was performed in longitudinal orientation (Figure 2E). Even in 2D scans we were able to observe the different orientations and geometries of RyR clusters, which were previously not discernible with techniques such as dSTORM or STED, predominantly due to their poorer axial resolution. Noteworthy among the clearly identifiable geometries were the end-on view of the dyads where the curvatures of the RyR clusters were visible (Figure 2F) and the side-on view of clusters located at transverse (Figure 2G) and longitudinal tubules (Figure 2H). In the latter views, a characteristic groove in the intensity topography of the RyR cluster allowed us to predict the geometry of the local t-tubule (dashed lines), even when fluorescent markers of the t-tubules were not available.

### Variable JPH2 sub-domains in peripheral and deeper couplons

10x EExM revealed punctate labelling morphologies of both JPH2 (cyan) and RyR (red) in the periphery of healthy rat ventricular myocytes. These RyR arrays were planar as reported previously (5) and therefore the mutual arrangement of RyR and JPH2 could be fully appreciated in 2D images similar to Figure 3A. The vast majority of the RyR clusters co-clustered with JPH2 (despite some exceptions, arrowhead), however the relative density of JPH2 across these domains varied from cluster to cluster. Figure 3B illustrates three exemplary clusters which consist of typically punctate patterns of RyR (dashed lines outline the RyR arrays), but also highly heterogeneous organisation of JPH2 within the cluster domain. Whilst JPH2 occupied the entire area in some clusters (Figure 3B-i) and only small sub-domains most others (Figure 3B-ii).

**Figure 3.**
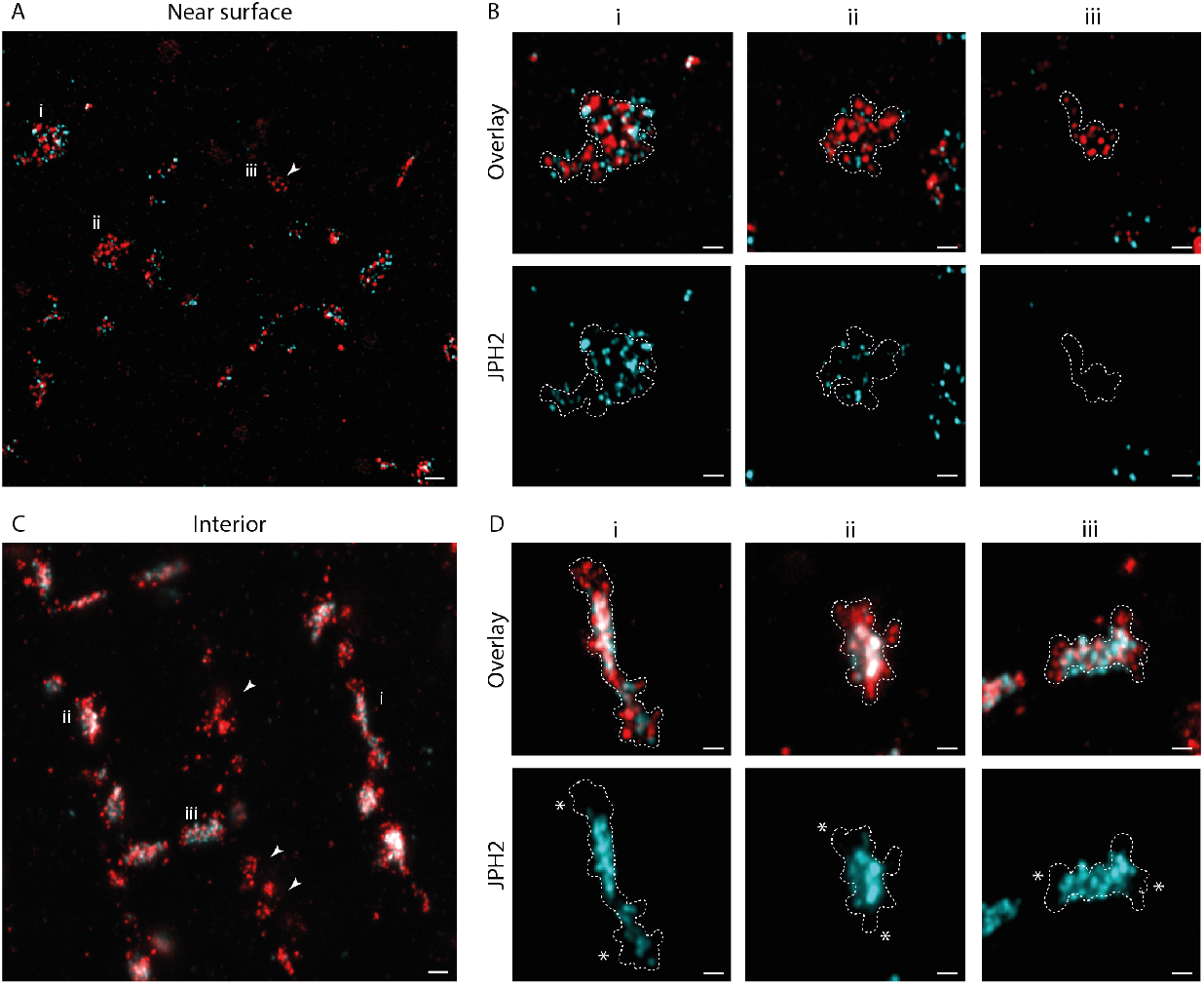
Molecular positioning of RyR and JPH2 near the cell surface and deeper in the cell interior. **(A)** A 10x EExM image taken near the surface of a ventricular myocyte illustrates the organisation of RyR2 clusters (red), co-localising frequently with punctate JPH2 labelling (cyan), across a 2D geometry of the nanodomain. A small proportion of (typically smaller) RyR clusters which lacked JPH2 were also noted (arrowhead). **(B)** Magnified view of three exemplar clusters (noted **i, ii** & **iii** in panel A) illustrate the boundary of the RyR cluster. JPH2 in these clusters show high variability in both the density and the spatial uniformity inside the RyR cluster boundary. **(C)** A shallow maximum-intensity projection of a 100-nm-deep 10x EExM image volume of RyR (red) and JPH2 (cyan) notably identifies more frequent RyR clusters devoid of co-localising JPH2 (arrowheads) in the cell interior. **(D)** Magnified views of three exemplar regions show that the RyR cluster boundaries (dashed lines) consistently extend beyond the longitudinal poles of the intrinsic JPH2 sub-domains (asterisks). Scale bars: A & C: 200 nm; B & D: 100 nm.

By comparison, RyR clusters devoid of JPH2 were more frequently observed in the interiors of these myocytes (arrowheads; Figure 3C). In clusters where RyR and JPH2 were co-localised, JPH2 appeared to be highly-enriched in clearly resolvable sub-domains (see exemplary clusters in Figure 3D-i, ii & iii). More notably, these sub-domains were located particularly in the middle portion of the RyR clusters, as viewed in side-on orientation. This meant that near the poles of the couplon (asterisks in Figure 3D), RyRs were less consistently co-localised with JPH2. A broadened shape of the RyR cluster (outline of the RyR cluster shown with dashed lines) and a looser organisation of the RyRs were also noted near the poles.

### RyR cluster fragmentation and molecular-scale fraying in RV failure

We applied 10x EExM to examine the remodelling of RyR cluster architecture in RV myocytes isolated from rats with MCT-induced RV failure (MCT-RV). We had previously used this approach to characterise the fragmented nature of the RyR clusters, along with a higher likelihood to be phosphorylated at Serine-2808 (5). Here, we sought to examine the accompanying changes to the structural component of the dyads relating to JPH2. Figures 4A & 4B compare exemplary 10x EExM images of RyR (red) and JPH2 (cyan) in the interiors of control RV (CON-RV) myocytes and MCT-RV myocytes respectively. At low magnification, both scenarios featured RyR clusters which were predominantly aligned transversely. In both cases, most of the RyRs appeared to co-cluster with JPH. The cluster fragmentation in MCT-RV cells featured distinct groupings of smaller RyR clusters in 1-2 μm-wide sub-cellular regions, which were otherwise devoid of larger RyR clusters (arrowheads in Figure 4A-right). Closer examination of these regions showed that each cluster fragment still featured smaller sub-domains of JPH2, however the RyR puncta appeared more scattered *around* the JPH2 sub-domain (Figure 4B-upper row). Examining the RyR position in relation to the JPH2 sub-domains (Figure 4B-lower row), the RyRs appeared to scatter around the edges of the JPH2 structural domains to a greater extent, resembling a fraying rope (see 3D isosurface visualisation in Figure 4C). The punctate nature of the JPH2 morphology in 10x EExM images allowed us to perform counts of the labelling units. From this analysis, we observed that there was a reduction in the JPH2 puncta in MCT-RV clusters in comparison to CON-RV (mean 5.88 JPHs per cluster in MCT-RV vs 10.65 in CON-RV; Figure 4D). This reduction was also accompanied by a ∼ 45% reduction in the mean density of JPH2 within the cluster volume (Figure 4E). Whilst we did not observe a statistically significant change in the mean ratio of the number of JPH2 puncta to the RyR puncta in a given cluster, we did observe a drop in the variability of this ratio in MCT-RV, compared to CON-RV (standard deviation 0.53 in MCT-RV vs 0.27 in CON-RV: 0.27; Figure 4F). The shift in JPH2:RyR ratio happened to a lesser extent for MCT-LV (Supplementary Figure 1).

**Figure 4.**
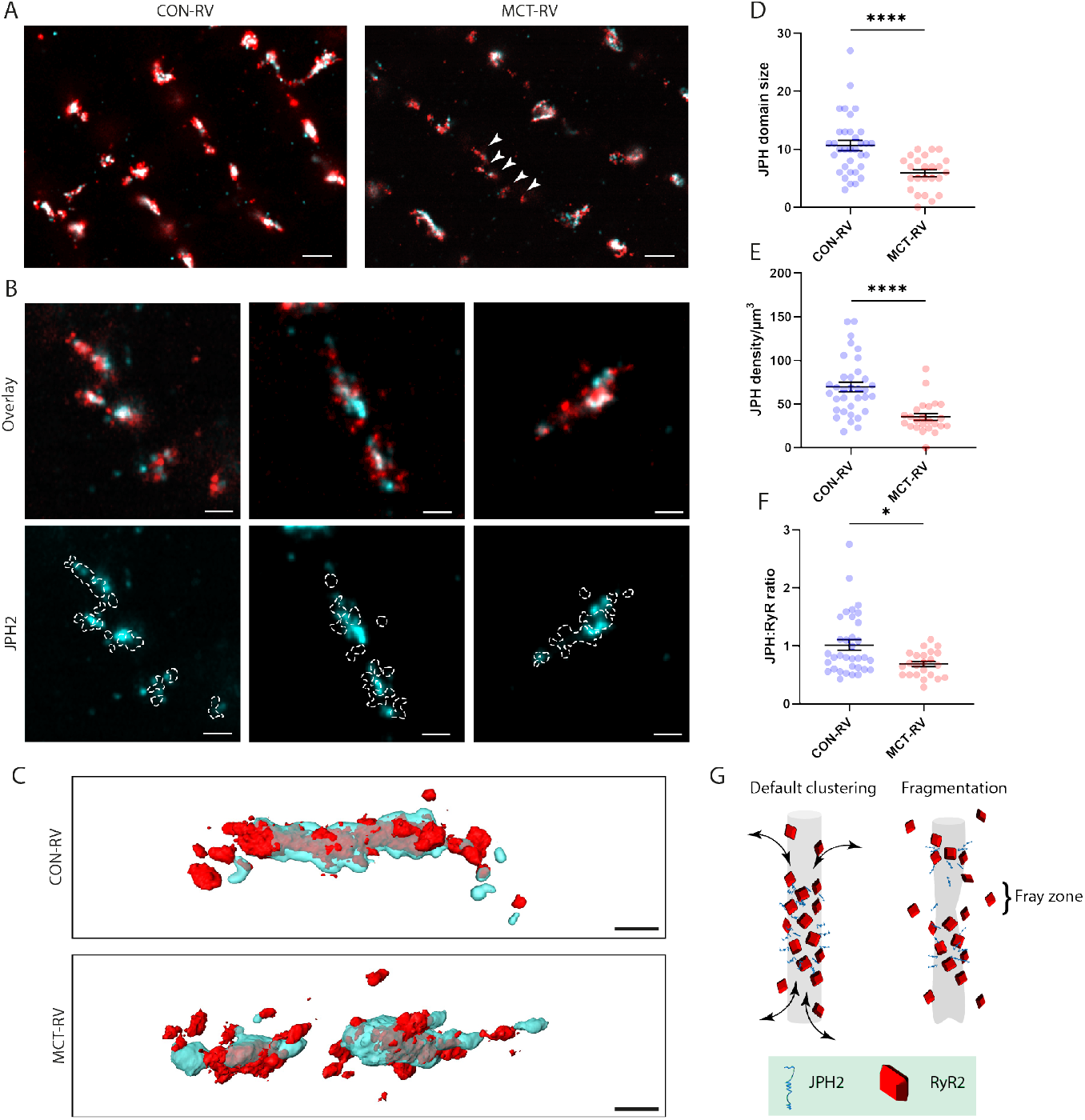
Molecular-scale remodelling of RyR clusters and intrinsic JPH2 organisation in RV failure. Exemplar 10x EExM images of RyR2 (red) and JPH2 (cyan) organisation in the interiors of myocytes isolated from CON-RV (left) and MCT-RV (right). Note the shorter RyR and JPH domains in the latter compared to the control (arrowheads). **(B)** Magnified view of three exemplar clusters (top row) show a fragmented RyR cluster morphology in MCT-RV (red) overlaid with the local JPH2 sub-domains (cyan). The bottom row depicts the poor alignment of the punctate RyR densities (*i*.*e*. frayed pattern) in relation to the JPH2 subdomains. **(C)** 3D Surface render of exemplar RyR clusters (red) and JPH2 sub-domains (cyan) from CON-RV (top) and MCT-RV (bottom) cardiomyocytes. Dot plots compare **(D)** the JPH2 domain size (in terms of detected JPH2 puncta per domain), **(E)** the density of JPH2 organisation within the nanodomain, **(F)** and the estimated ratio of JPH2:RyR ratio between CON-RV and MCT-RV. **(G)** Schematic depicting hypothesized sub-domain remodelling in RyR clusters. In the healthy phenotype, JPH2 occupies a central sub-domain of the dyad whilst the poles, devoid of JPH2, may serve as turnover domains (arrows) where RyR organisation may be either looser or physically broader. In maladaptive remodelling, the downregulation of JPH2 may lead to fragmentation of the structural sub-domain along with potential remodelling of the local t-tubule and concurrent fragmentation of the RyR cluster, in an unknown temporal sequence. The fragmentation of the JPH2 sub-domain may cause “fray zones” which function as turnover domains, giving rise to the dissipated or frayed morphology observed in EExM image data. Mann-Whitney tests (****: p < 0.0001, df = 57; *: p = 0.0164, df = 56). Whiskers notate mean and standard error. Scale bars: A: 500 nm; B & C: 100 nm.

## Discussion

### EExM as a tool for visualising topologies and molecular organisation of dyads

A handful of optical super-resolution techniques, including DNA-PAINT and EExM (5), currently offer the capability to resolve individual RyRs, *in situ*. However, a major shortcoming in the use of localization-microscopy to visualise cardiac dyads has been the limited capacity to resolve 3D complexity of the geometries and the topological features. We previously predicted the possibility of fully-resolving individual RyRs within the curvatures of the dyad (see a simulation in the supplementary figure S5 of (5)). The 3D visualisations shown in this paper demonstrate the full range of geometries and morphologies of RyR clusters observed in cardiomyocytes using 10x EExM. 4x EExM provides an arguably more versatile mode of imaging (particularly compatible with cryosections for imaging the transverse aspect of myocytes), with visualisation of RyR clusters comparable to that achievable with 3D dSTORM. We observe that a complete visualisation of the tessellation of RyR clusters with the extensive t-tubule network that extend throughout the z-line can be achieved by imaging myocytes that are sectioned in a transverse orientation.

The most significant advantage of using EExM for 2D or 3D imaging of cardiac RyR clusters is that it can report the precise boundaries of the cluster, the likely geometry of the dyad as well as the positions of the RyRs. By comparison, dSTORM (in a typical 2D- or 3D-localisation implementation) lacks the localisation precision to localise individual channels. Despite reporting clear outlines of the clusters, the morphology within that outline was often spurious, as demonstrated before (21). The advantage in choosing techniques with sub-15 nm in-plane resolution (and ideally similar axial resolution) is that observing protein cluster features such as topology, co-localisation measurements and molecular-scale remodelling become more straightforward to observe. On this principle, more details can be gained by 10x or higher-order ExM implementations, such as adaptations of iterative ExM (33). However, pragmatically a balance needs to be sought between the resolution gains and the ease of the chosen expansion protocol; with the current state of the art, we find that 10x EExM is a convenient protocol which works well with isolated cardiomyocytes.

### Sub-domains of JPH2 occupancy and model of dyad fraying in pathology

We have observed distinct sub-domains of JPH2 within the dyad, which have not been observed with previous dSTORM imaging due to insufficient resolution. In sub-sarcolemmal couplons, the organisation of JPH2 may be more variable, leading to a structure which is more conducive to the rapid remodelling or dynamic re-organisation seen previously (30,34). In dyads located deeper into the cell interior, JPH2 localises more consistently to structural sub-domains occupying the central region. The looser co-clustering of RyR with JPH2 at the poles of the dyads may represent turnover domains which could enable exchange of RyRs (similar to turnover of gap junction plaques (35)). Such a polarity of RyR organisation in the dyads may indicate the local connectivity of the SR/ER which would form trafficking routes and lateral insertion points for dyad proteins, similar to those seen in synaptic proteins (36). The denser occupation of JPH2 in the central portions of the larger RyR clusters could indicate a lower propensity for remodelling or mobility of these domains.

In MCT-induced RV failure, we have consistently observed RyR cluster remodelling, particularly fragmentation of clusters, which coincides numerous maladaptations such as downregulation of structural proteins such as BIN1 & JPH2 (either *via* microRNA silencing, calpain cleavage or trafficking misdirection) (22,37). By quantifying JPH puncta and labelling density we have demonstrated that there is JPH2 downregulation inside the MCT-RV dyad geometry (an observation made previously by western blot (37)). Our spatially-resolved analysis shows that the JPH2:RyR ratio becomes less heterogeneous in MCT-RV. This may support the hypothesis that there are un-coupled pools of JPH2 resident within each dyad (21), capable of buffering a global downregulation in JPH2 expression. Such a depletion could be seen in our approach as a change in the local co-clustering ratio of JPH2 and RyR.

In MCT-RV myocytes, the JPH2 sub-domains followed the fragmented phenotype observed in the RyR distribution, supporting the idea that regressing and redistributing structural proteins have a role in the intrinsic dyad and t-tubular remodelling. For example, the appearance of RyR dispersion in ‘fray zones’, in close proximity to regions where JPH2 sub-domains have either regressed or fragmented, could result from one or more possible maladaptations in the maintenance of the dyad. Firstly, it is possible that the regression of the JPH2 sub-domain in a formerly-mid region of a dyad creates new turnover domains through which RyRs can be mobilised (Figure 4G). Alternatively, the fray zones may indicate focal points where the labile membrane topologies which make up the t-tubules undergo either pinocytosis or new branching (38).

### Bottlenecks and motivations for expansion microscopy of cardiac RyR clusters

EExM, despite the far superior 3D resolution that it offers in resolving both the topologies of the dyads and individual RyRs, still requires a number of calibrations and tests. These include independent estimates of the gel expansion factors and isotropy of expansion (see detailed discussion in (23)). Combining the X10 expansion protocol with imaging modalities such as STED or SIM can further improve the resolution attainable to a sub-10-nm regime. The skill and experience involved in the handling and fluorescence imaging of the hydrogels remain a significant human element in the methodology. However, the emergence of newer gel formulae capable of greater expansion isotropy (39), and pan-stains which can delineate other cellular compartments (40), enable increasingly greater reliability and versatility of ExM for investigations such as this. Despite the unique opportunity to visualise the transverse aspect of the t-tubular network and dyads with 4x EExM combined with immunohistochemistry, we note the excessive rigidity (due to the extracellular matrix) inherent to myocardial tissue sustaining pathological remodelling. Whilst the size and specificity of the fluorescent probes are critical (the RyR2 and JPH2 antibodies used here have been extensively characterised previously), it should also be noted that post-labelling and re-embedding ExM gels can further improve the completeness and geometric accuracy of the structures (41).

The far superior 3D information offered by EExM compared to other recent implementations of super-resolution microscopy remains a key reason for its utility. The 3D contextual information it has provided us in terms of t-tubule geometries, organisation of structural proteins such as JPH2, and examining RyR cluster fragmentation in heart failure has been pivotal to our observation of JPH2 sub-domains. Having used this method in our recent studies, we also observe the highly accessible nature of the EExM protocols compared to other super-resolution techniques. We therefore anticipate broad uptake and refinement of the method within the cellular cardiac discipline in the next few years.

## Conclusions

We have demonstrated that combining EExM with 3D visualisation can reveal the diverse topologies and geometries of RyR clusters in cardiac muscle. Owing to the fine in-plane and axial resolution afforded by 10x EExM, we have been able to identify sub-domains within the cardiac dyad which are occupied by the structural protein JPH2. In RV failure, we observe fragmentation of these JPH2 sub-domains, coinciding with a drop in the intrinsic JPH2 density and the dyad-to-dyad heterogeneity in the ratio between RyR and JPH2. Using 3D visualisation of individual RyRs, we identify waning or fragmenting JPH2 sub-domains as the foci of RyR channel dispersion and likely sites of t-tubule remodelling.

## Acknowledgements

The authors acknowledge research funding from the DiMeN doctoral training programme of the Medical Research Council, UK Research and Innovation (grant no. MR/S03241X/1) and the Wellcome Trust (207684/Z/17/Z). We acknowledge Dr S Boxall and the University of Leeds bio-imaging facility for technical assistance. We also thank Dr Michael Colman (University of Leeds) for helpful discussions.

## Author contributions

IJ, EW & JC conceived experiments. IJ, AJS, JC provided supervision. TMS, MH performed isolation experiments. TMS performed imaging and made primary observations. TMS & IJ performed image analysis. TMS & IJ wrote the manuscript.

## Materials and Methods

### Animal model

Experiments were performed according to the UK Animals (Scientific Procedures) Act of 1986 and with UK home office approval and local ethical approval. Animals were housed at 20-22°C, 50% humidity, in a 12-hour light/dark cycle and were given *ad libitum* access to food and water. Adult male Wistar rats weighing 200 ± 20 g were given a single intraperitoneal injection of 60 mg/kg (of body weight) crotaline (Sigma-Aldrich) (dissolved in HCl and 140 mM NaCl, pH 7.4) to induce pulmonary arterial hypertension, as detailed previously (22). Control animals were injected with an equivalent volume of saline solution (140 mM NaCl). Animals were weighed 3 times weekly for the first 3 weeks, then daily near the heart failure period (between days 21 and 28). When signs of heart failure were observed (piloerection, cold extremities, lethargy, dyspnoea, consecutive days of weight loss or a 10 g weight loss in a single day) animals were humanely euthanised by concussion followed by cervical dislocation. Control animals were taken on the median survival day of failing animals.

### Cardiomyocyte isolation

Individual cardiomyocytes were isolated from freshly dissected hearts by the Langendorff perfusion method, as detailed previously (5). The ventricles were separated prior to shaking to yield separate cell suspensions. Cardiomyocytes were attached to imaging chambers (laminin-coated coverslips attached to acrylic slides) for 2 hours at 37°C and fixed in 2% paraformaldehyde (Sigma-Aldrich) (w/v) in phosphate-buffered saline (PBS, Sigma-Aldrich) at room temperature for 10 minutes. Samples were stored in fixed cell storage solution containing 0.1% sodium azide (NaN_3,_ Sigma-Aldrich) and 5% bovine serum albumin (BSA, Thermo Fisher Scientific), until immunofluorescent labelling.

### Immunofluorescence labelling and antibodies

Cells were permeabilised with 0.1% Triton X-100 (Sigma-Aldrich) in PBS for ten minutes and then blocked with 10% normal goal serum (NGS, Thermo Fisher Scientific) in PBS for one hour. Samples were incubated with primary antibodies (RyR2: MA3-916; JPH2: 40-5300, Thermo Fisher Scientific) overnight at 4°C, diluted to 1:200 and 1:250 respectively in incubation solution containing (w/v or v/v) 0.05% NaN_3_, 2% BSA, 2% NGS and 0.05% Triton X-100 dissolved in PBS. Samples were washed in PBS and then incubated with secondary antibodies (anti-mouse Alexa Fluor 488, anti-rabbit Alexa Fluor 594, Thermo Fisher Scientific) for two hours at room temperature, diluted to 1:200 in incubation solution. Samples were washed in PBS and imaged to obtain pre-expansion images.

### Cardiac tissue processing and immunohistochemistry

For immunohistochemistry, cardiac tissue sections were obtained according to the following protocol. Following dissection into the respective ventricles, cardiac tissue was fixed in 1% paraformaldehyde (w/v) in PBS at 4°C for 1 hour. Fixed tissue was then washed and cryoprotected in sucrose solution (series of 10%-20%-30%). Excess sucrose solution was removed before a thin layer of OCT was applied to coat the tissue. The tissue was snap frozen for 2 minutes by immersion in methylbutane within a container of liquid nitrogen. Frozen tissue blocks were cryosectioned with a Feather Blade at −20°C. 10 µm-thick sections were obtained and were attached to coverslips until immunofluorescent labelling.

Cardiac tissue sections were treated with Image-iT FX signal enhancer (Thermo Fisher Scientific) for 1 hour at room temperature prior to incubation with primary antibodies (RyR2: MA3-916 (Thermo Fisher Scientific); NCX: R3F1 (Swant); Cav3: 610420 (BD Transduction), overnight at 4°C, and diluted 1:200 in incubation solution. After washing in PBS, sections were then incubated with secondary antibodies (as above) for two hours at room temperature, in incubation solution.

### Expansion microscopy

First, samples were incubated with 0.1 mg/ml acryloyl-X (Thermo Fisher Scientific) in PBS overnight at 4°C, then washed in PBS immediately prior to addition of gel solution.

We performed 10x EExM on isolated cardiomyocytes as detailed previously (5,23). X10 gels (4:1 molar ratio of dimethylacrylamide (Sigma-Aldrich) and sodium acrylate (Sigma-Aldrich), dissolved in deionized H20 (dH_2_O)) were made according to the previous recipe (24,25). Gel solution was made fresh and bubbled with nitrogen gas for one hour on ice. Potassium persulfate (Sigma-Aldrich) was added from a fresh 0.036 g/ml stock to 0.4% molar relative to the monomer concentration and the solution was bubbled for another 15 mins on ice. 500 µl of the gel solution was mixed rapidly with 2 µl of N,N,N’,N’-Tetramethylethylenediamine (Sigma-Aldrich) and quickly added to the sample coverslip. The polymerisation chamber, comprising the sample coverslip with two coverslip spacers either side, was sealed with a top coverslip. Gels were polymerised after two hours. The major axes of the gel were measured to calculate the pre-expansion size.

We also performed 4x EExM on tissue sections, similar to that described for isolated cardiomyocytes previously (23). The gel solution, according to the proExM protocol previously described (26), was made in advance and defrosted from frozen aliquots. Tissue sections were incubated with monomer solution containing (w/v, Sigma-Aldrich) 8.6% sodium acrylate, 2.5 % acrylamide; 0.15% N,N’-Methylenebisacrylamide, 11.7% NaCl, PBS, 0.1% ammonium persulfate (APS) and N,N,N’,N’-Tetramethylethylenediamine first for 30 mins at 4°C and then for 2 hours at 37°C.

Polymerised gels were removed from the coverslip chamber and placed into 6-well plates to undergo digestion in 0.2 mg/ml proteinase K (New England Biolabs) dissolved in digestion buffer (50 mM Tris pH 8.0 (Thermo Fisher Scientific), 1mM ethylenediaminetetraacetic acid (Sigma-Aldrich), 0.5% Triton X-100, 0.8M guanidine HCl (Sigma-Aldrich) and dH_2_0) overnight at room temperature. Gels were expanded by shaking in dH_2_0 until the gel expansion reached a plateau, replacing the dH_2_0 every hour. The final gel size was measured to calculate the macroscale expansion factor, in relation to the pre-expansion size.

### Image acquisition

Expanded gels were placed into acrylic chambers with a square cut-out, attached to a no. 1.5 glass coverslip (Menzel Gläser), which had been coated with 0.1% (v/v) poly-L-lysine (Sigma-Aldrich) at room temperature for 30 minutes. Airyscan imaging was performed on an inverted LSM880 (Carl Zeiss, Jena), with a Plan-Aprochromat 63x 1.4 NA objective with a working distance of 0.19 mm. Alexa Fluor 488 and Alexa Fluor 594 were excited with 488 nm and 561 nm DPSS lasers, while emission bands were selected using the in-built spectral detector. Pixel sampling of primary data was ∼40 nm per pixel in x/y, ∼180 nm in z. Airyscan datasets were subjected to a pixel reassignment and a linear Wiener deconvolution *via* the Zen software.

### Image analysis

3D count and distance analysis of RyR and JPH2 was performed in Fiji using the plugin ImageJ3D suite (27). Individual clusters were cropped and background subtracted (following gaussian blur). A 3D iterative thresholding algorithm separated objects (relating to individual RyR and JPH2 puncta). A custom-written Python code (using pandas and openpyxl libraries) was used to count the number of RyR and JPH2 per cluster. All scale bars shown in this paper indicate the original spatial scale of the sample; in EExM images, the scale shown is corrected by the sample’s estimated expansion factor, determined by the local sarcomere length, as previously established (5).

---

**Supplementary Figure S1.**
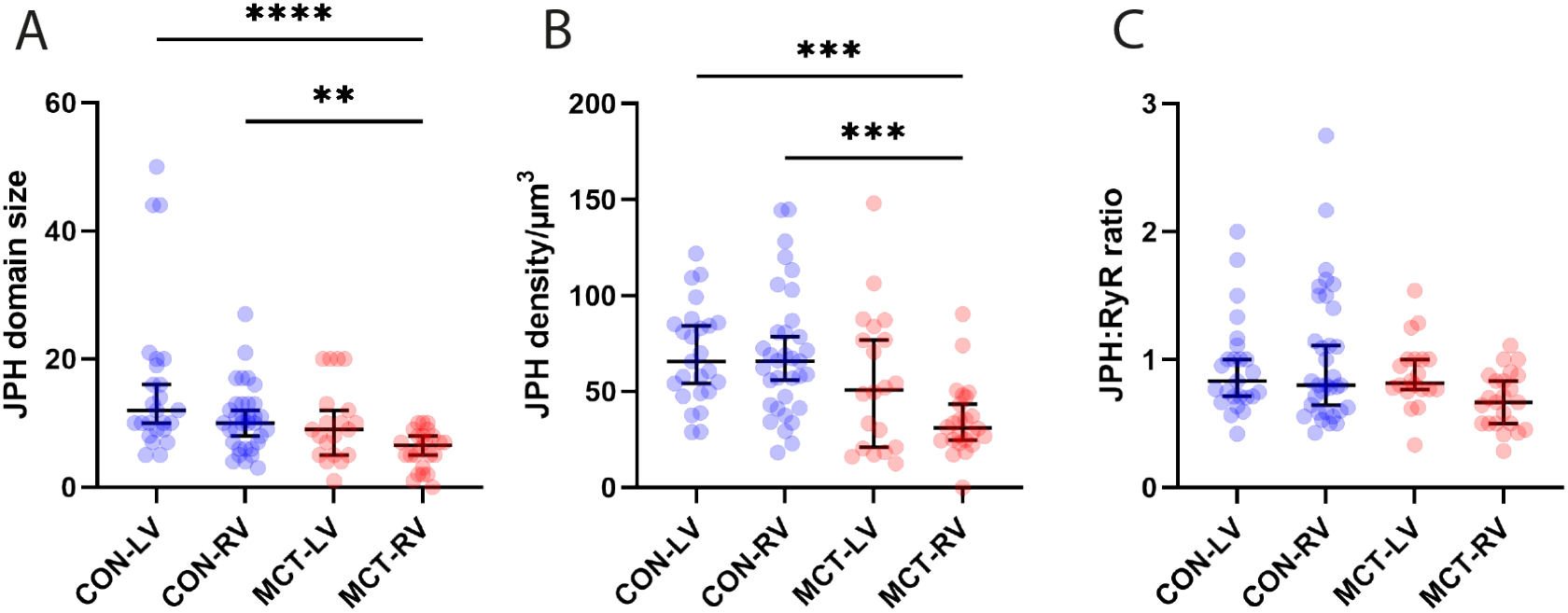
Four-chamber comparison of RyR and JPH remodelling in RV failure. Dot plots compare **(A)** the JPH2 domain size (in terms of detected JPH2 puncta per domain), **(B)** the density of JPH2 organisation within the nanodomain, **(C)** and the estimated ratio of JPH2:RyR ratio, between CON-LV, CON-RV, MCT-LV and MCT-RV. Kruskall-Wallis tests (****: p < 0.0001, df = 102; ***: p < 0.001, df = 102; **: p = 0.0016, df = 101). Whiskers notate median and 95% confidence interval.

